# Antidepressant Sertraline Hydrochloride Inhibits the Growth of HER2+ AU565 Breast Cancer Cell Line through Induction of Apoptosis, and Arrest of Cell Cycle

**DOI:** 10.1101/2021.09.07.459321

**Authors:** Atia-tul- Wahab, Sharmeen Fayyaz, Rimsha Irshad, Rafat A. Siddiqui, Atta-ur- Rahman, M. Iqbal Choudhary

**Affiliations:** Dr. Panjwani Center for Molecular Medicine and Drug Research, International Center for Chemical and Biological Sciences, University of Karachi, Karachi-75270, Pakistan; H. E. J. Research Institute of Chemistry, International Center for Chemical and Biological Sciences, University of Karachi, Karachi-75270, Pakistan; Food Chemistry and Nutrition Science Laboratory, College of Agriculture, Virginia State University, VA 23806, USA; Department of Biochemistry, Faculty of Science, King Abdulaziz University, Jeddah-21589, Saudi Arabia

**Keywords:** Sertraline hydrochloride, Breast cancer, Drug repurposing, Selective serotonin re-uptake inhibitors

## Abstract

Breast cancer is one the most aggressive cancer worldwide, especially Pakistan due to limited therapeutic options. This study was conducted to repurpose the use of selective serotonin reuptake inhibitors (SSRIs), in the treatment of breast cancers, and merit to pursue drug re-positioning in oncology. Anti-proliferative activity of SSRIs, such as fluoxetine, paroxetine, and sertraline hydrochloride on the growth of AU-565, MCF-7, MDA-MB-231, and BT-474 breast cancer cell lines, along with human fibroblast BJ cells was determined *in vitro*. Changes in nuclear morphology (DAPI staining), and induction of apoptosis (flow cytometry, and caspase-3 activation) were also studied. Sertraline hydrochloride most effectively inhibited the growth of breast cancer cells *in vitro.* Therefore, pharmacological mechanism involved in sertraline mediated cell death was investigated in HER2+ AU565 cell line. Enhanced nuclear fragmentation, increased Annexin (+) cells, and caspase-3/7 activation indicated that sertraline-mediated cell death could be a result of BCl2-independent apoptosis as evidenced by expression of Bax, and BCl2 genes. Taken together, our results identified sertraline hydrochloride, as a potential candidate for the treatment of HER2-positive breast cancer. Even though these are *in vitro* results, this study opens great opportunity in the field of drug repurposing for the development of chemotherapeutic agents.

## 1. Introduction

Breast cancer is the second leading cause of morbidity and mortality in women. Over 250,000 cases of invasive breast cancers are diagnosed in the United States [1] and 40,000 patients die from this disease every year [2]. According to a recently published report, Pakistan has the highest incidence of breast cancer in Asia. According to an estimate, 83,000 cases are being annually reported in the country, and over 40,000 deaths are caused by it [3, 4].

Breast cancers are classified based on the expression of certain biomarkers, such as estrogen receptor (ER), progesterone receptor (PR), and human epidermal growth factors (HER2). Oncogene HER2 is overexpressed in approximately 15-20% of breast cancer cases. Poor prognosis, drug resistance, relapse, and metastasis are usually associated with HER2 overexpressed breast cancers [5]. For the last two decades, study of surface receptor HER2 has been of much interest because treatment with antibodies, such as trastuzumab, and pertuzumab, have improved the survival rate upto 30-40% [6]. Although the use of monoclonal antibodies accomplished excessive outcomes, the majority of patients still become resistant to therapy, also the disease relapse is also a common problem associated with the use of antibodies. The incidence of central nervous system metastases in patients with metastatic breast cancer who received therapy with trastuzumab in first line is in the range of 41 – 48%. Thus, drug resistance, and relapse are important clinical challenges, which necessitate the development of new and innovative approaches to overcome the resistance of current anti-cancer drugs [7].

Both clinical, and experimental studies have demonstrated that depression has an influence on tumor progression [8, 9]. National Institute of Aging provided compelling data that chronically depressed people over the age of 70 are 88% more likely to develop cancer [10]. Moreover, depression is also a common problem of cancer patients, 25 - 58% of cancer patients are at risk of developing depression [11]. Co-existing depression have negative impact on cancer treatment, and results in decreased survival in breast cancer patients [12].

There has been a rapid increase in the use of antidepressant drugs in the last decade. Women are twice likely users of antidepressants than men. A recent Indian survey showed that a total of 62.2% patients were using selective serotonin reuptake inhibitors (SSRIs) [13]. SSRIs have also been the most commonly prescribed antidepressants in America. Annual prescriptions for sertraline have increased dramatically from 10.8 million in 2006 to 35.7 million in 2010 [14].

SSRIs is an important class of anti-depressants, and frequently prescribed for the treatment of depression, obsessive–compulsive disorder, and other related disorders [15]. SSRIs act as reuptake inhibitor of the neurotransmitter serotonin, by blocking the action of the serotonin transporter. This in turn increased the extracellular level of serotonin, a neurotransmitter, which is often referred as “feel good hormone” [16].

Interestingly, some antipsychotic drugs, such as trifluoperazine, chlorpromazine, fluphenazine, and aripiprazole, have also shown cytotoxicity against lung cancer cell lines [17]. Goyette *et al*. also reported that members of phenothiazine class, used as dopamine receptor antagonists (anti-psychotic), potently inhibited the growth, and proliferation of metastatic breast cancer cell line MDA-MB-231 [18].

Rosetti, and co-workers have reported that paroxetine, a serotonin reuptake inhibitor, induced a marked cytotoxic effect in various human and murine cell lines [19]. Reduced risk of colorectal cancer was observed in patients with a daily intake of SSRIs, as compared to tricyclic anti-depressants [20]. Caspase mediated apoptosis is reported in Burkitt lymphoma cells in the presence of paroxetine, fluoxetine, and citalopram [21]. These studies showed the oncolytic effect of anti-depressants, specifically SSRIs. It is, therefore, important to further study the beneficial effects of SSRIs in other cancer models.

The aim of the present study was to study the effect of SSRIs, paroxetine, fluoxetine, and sertraline hydrochloride (Fig 1) on different breast cancer cell lines, and human fibroblast cell line. Our results indicate that among all the drugs evaluated, sertraline hydrochloride, most effectively inhibited the growth and viability of breast cancer cells *in vitro*. Therefore, the possible mechanisms underlying the cytotoxic effect of sertraline hydrochloride was also studied. We observed that sertraline hydrochloride has induced apoptosis, and arrested cells in the G2/M phase of cell cycle. Our findings, therefore, explore the possibility of repurposing sertraline hydrochloride for the treatment of breast cancer.

**Fig 1.**
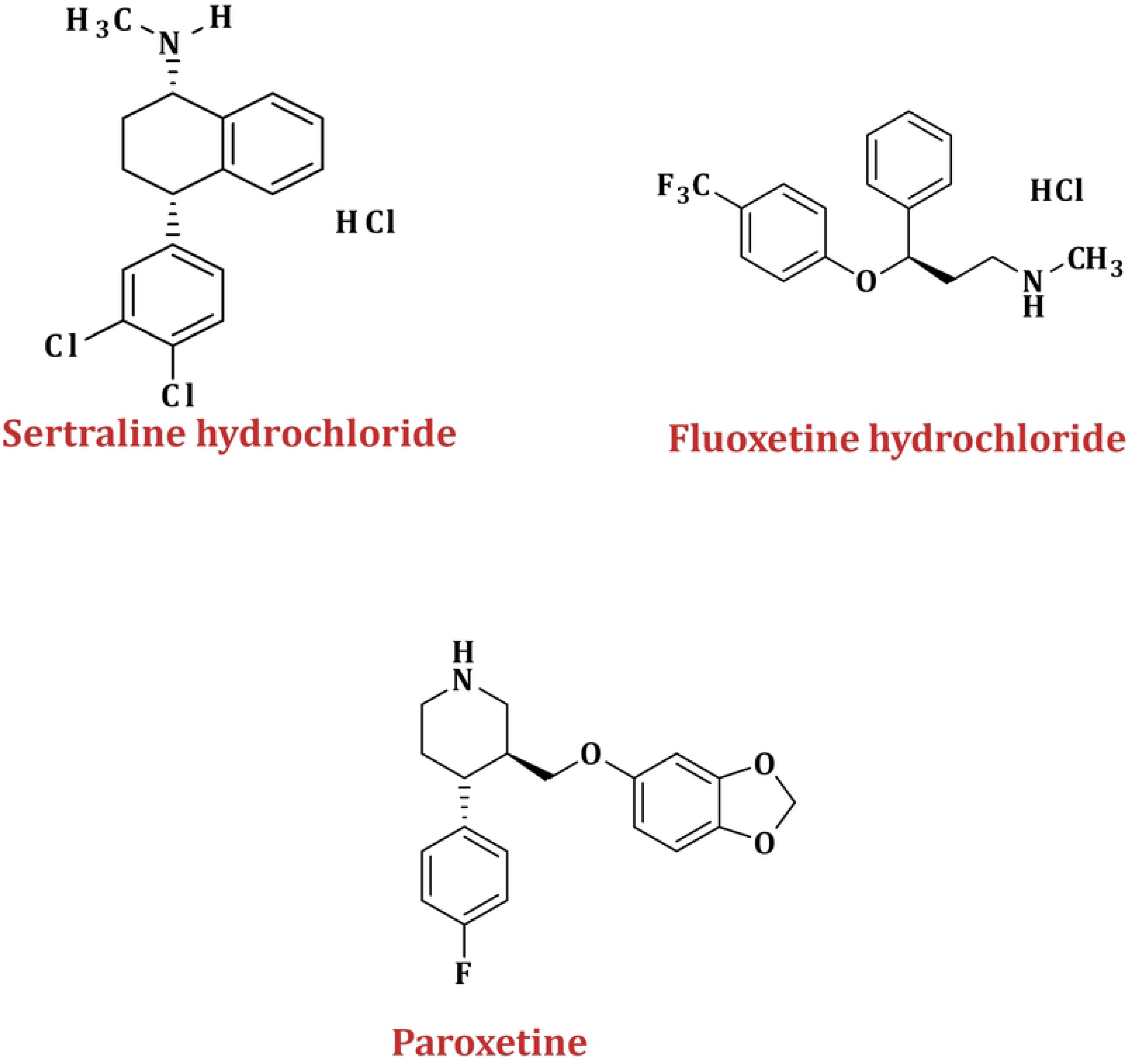
Structures of anti-depressant SSRIs used in the study

## 2. Materials and Methods

All drugs were obtained from Drug Bank of PCMD (ICCBS), University of Karachi, Pakistan. Doxorubicin was purchased from Sigma Aldrich (Merck KGaA, Germany). Propidium iodide (Biosera, France), 4′,6-diamidino-2-phenylindole (DAPI), and annexin V-FITC (Life Technologies, US). cDNA synthesis kit (cat no. K 1622, Molecular Biology, USA), and SYBER Green Master Mix (cat no. K 0221 Thermo Fischer Scientific, USA) were used. Primers for real time PCR were purchased from Macrogen, Inc., South Korea.

### 2.1 Cell Culture and Maintenance

Breast cancer cell line AU565 (ATCC CRL-2351) was purchased from the American Type Culture Collection (ATCC), and maintained in the ATCC modified RPMI media. Growth medium was supplemented with 10% fetal bovine serum, 1% L-glutamine, and 1% sodium pyruvate. All the chemicals were purchased from Gibco, US. Cells were sub-cultured twice a week.

### 2.2 Cell Viability

Cell viability was determined using colorimetric 3-(4,5-dimethylthiazole-2-yl)-2,5-diphenyl tetrazolium bromide (MTT) reagent. Briefly, cells were seeded at a density of 1 x 10^4^ cells/well (100 μL) in a 96-well plate. After 24 h, cells were treated with the desired concentrations of different anti-depressants (fluoxetine hydrochloride, sertraline hydrochloride, and paroxetine hydrochloride), and incubated for 48 h. MTT solution (5 mg/mL in PBS) was added to each well, and left for a second incubation of 3-4 h. The supernatant was aspirated, and formazan crystals were dissolved in 100 μL DMSO. Absorbance was recorded at 550 nm with a microplate reader (Molecular Devices, USA).

### 2.3 Annexin V-FITC/PI Double Staining Assay

Apoptotic or necrotic cell death was analyzed by dual staining of cells with FITC-labelled Annexin V, and propidium iodide (PI). Cells were harvested after treatment of cells with different concentrations of sertraline hydrochloride for 48 h. Both adherent, and floating (live and dead) cells were collected, centrifuged at 2,500 rpm, and washed with ice-cold buffer. The washed pellet was re-suspended in annexin binding buffer (500 μL), and stained with annexin-V, and PI using 5 μL, and 1 μL, respectively. Samples were incubated in the dark for 15 minutes before analysis [22]. Induction of apoptosis was determined using BD FACS Caliber Cell Flow Cytometer using CellQuest software (BD Bioscience, USA).

### 2.4 Cell Cycle Analysis

The effect of sertraline hydrochloride on cell cycle progression was analyzed by Fluorescence Activated Cell Sorter (FACS) of 10,000 cells, stained with propidium iodide (PI). Cells were cultured in 25 cc flasks, and treated with varying concentrations (0, 7, 14, and 28 μM) of sertraline hydrochloride for 48 h. Cells were then harvested, and fixed in 500 μL of chilled 70% ethanol. Cell pellet was washed twice with ice cold PBS (at 300x g), and incubated in the dark for 30 min after addition of 100 μg/mL Rnase A and 50 μg/mL of PI (125 μL, each) at room temperature. DNA content was examined based on the PI signal using flow cytometry, and percentage (%) of cells in each phase was determined by using Flowjo software (BD Life Sciences, USA).

### 2.5 Gene Expression Analysis using RT-PCR

Total RNA of cells was extracted after treatment of cells with sertraline hydrochloride for 72 h using TRIzol reagent (Cat no. 15596-018, Ambion, Life Technologies, USA). Complementary DNA (cDNA) was prepared with 1 μg of RNA using revertaid cDNA synthesis kit (cat no. K 1622, Molecular Biology, USA), following manufacturer’s protocol. For quantitative real time PCR, cDNA was amplified with SYBR Green and ABI StepOnePlus Real-time PCR Systems. The relative expression was normalized to GAPDH expression, and was calculated by the 2^ΔΔCt^ method. Primer sequences are given in Table 1.

**Table 1:**
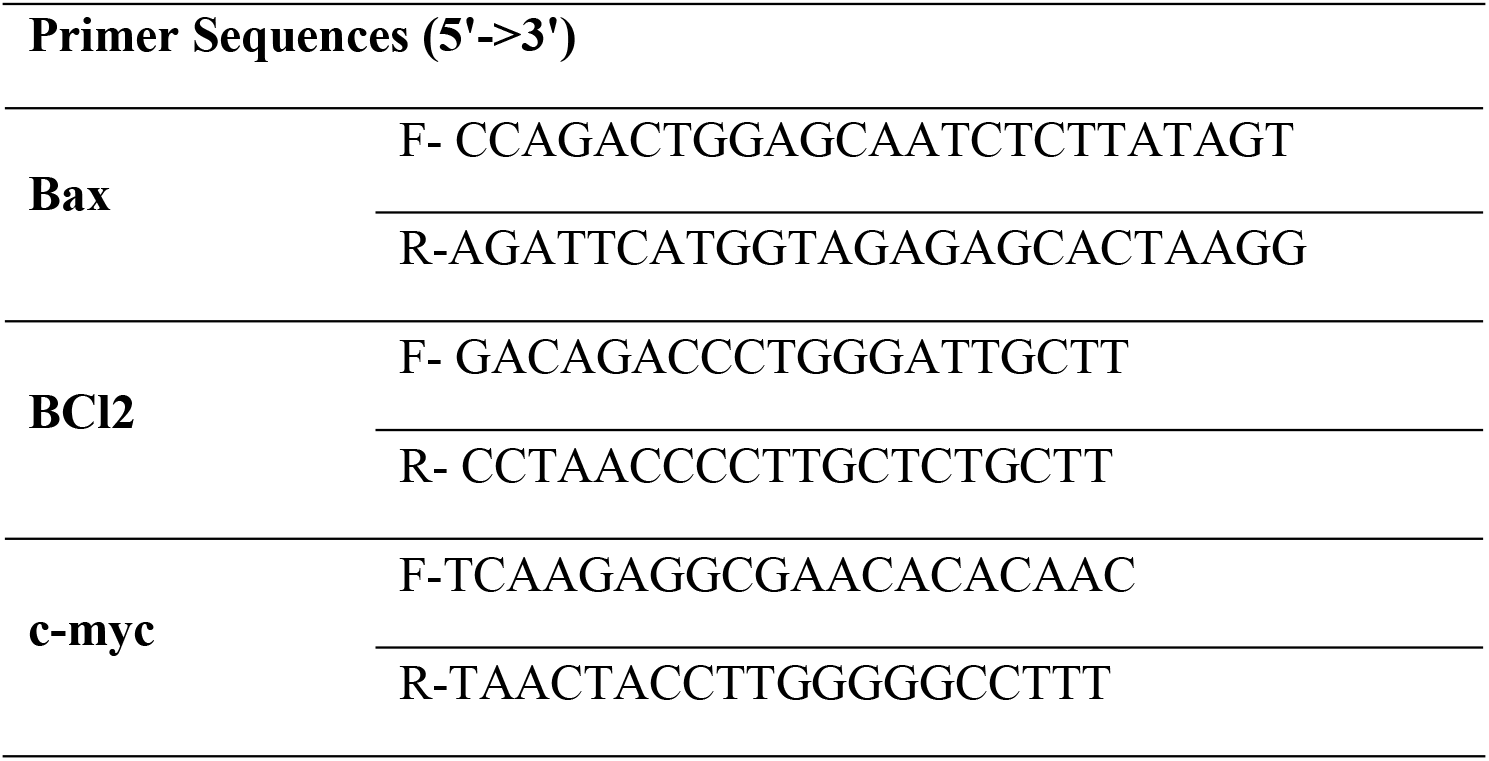
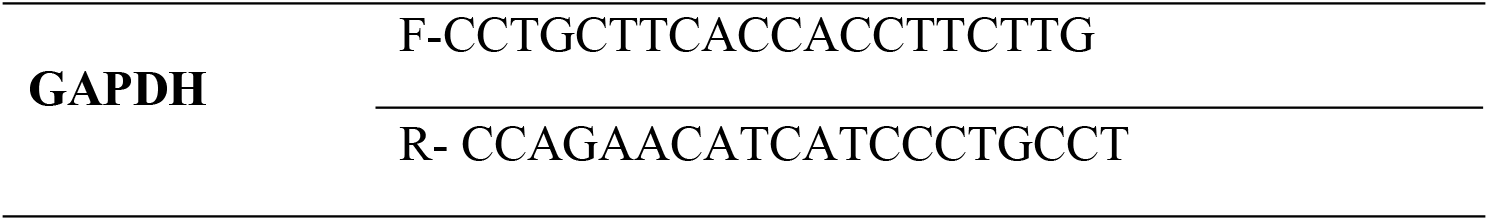
Primer sequences used for semi-quantitative real time PCR.

### 2.6 DAPI Staining

Induction of apoptosis was further confirmed by analysis of morphological changes in cell nuclei by using DAPI staining. AU565 cells were cultured in 24-well plates, and treated with different concentrations of sertraline hydrochloride for 48 h. Exhausted media was then removed by washing cells twice with PBS, and cells were fixed using 4% paraformaldehyde for 1 h. After fixation, cells were stained with DAPI stain (0.1 ng/0.5 mL) for 10 min at room temperature in the dark. Unbound DAPI stain was removed by washing the cells with PBS. Cells were analyzed under the microscope (TE-2000, Nikon, Japan) at 20X for morphological changes.

### 2.7 Caspase-3/7 Activity Analysis

Caspase-3/7 activity was quantified by fluorometric assay using ApoONE Homogeneous Caspase-3/7 Assay Kit (Promega, US). AU565 cells, after treatment with varying concentrations of sertraline hydrochloride for 24 or 48 h, were washed twice with PBS to remove media traces, and then incubated with 60 μL Apo-ONE lysis buffer for 1 h at room temperature. Caspase-3, and −7 activity was measured using Apo-ONE caspase-3/7 reagent. Fluorescence was measured continuously for 4 h with an interval of 30 min using Varioskan Lux multimode reader (Thermo Fisher Scientific, USA), with an excitation/emission wavelength of 495/532 nm. All measurements were carried out in triplicate, as reported by Sturzu, *et al.*, 2016 [23].

### 2.8 Statistical Analysis

Data is represented as mean ± SD. IC_50_ values were calculated using EZ-FIT software by Perrella Scientific, Inc., NH, USA. Significant differences between treated and control groups were analyzed by using one-way ANOVA with Tukey’s test as a post-hoc comparisons using Graphpad Prism software (GraphPad, CA, USA). Differences were considered statistically significant at *p* < 0.05.

## 3. Results

### 1.1 Viability of Breast Cancer Cells Decreased after Treatment with Anti-depressant Drugs

Paroxetine, and sertraline hydrochloride moderately inhibited the growth of breast cancer cell lines (Table 2) with IC_50_ values 13.8 ± 1.1 μM (AU565), 17.54 ± 1.6 μM (BT-474), 14.6 ± 0.57 μM (MCF-7), 9.4 ± 1.2 μM (MDA-MB-231) for sertraline hydrochloride. IC_50_ Values for paroxetine hydrochloride were 18.4 ± 0.63 μM (AU565), 15.67 ± 1.9 μM (BT-474), 16.18 ± 6.2 μM (MCF-7), and 13.56 ± 0.42 μM (MDA-MB-231). Fluoxetine hydrochloride weakly inhibited the breast cancer cells *in vitro* with IC_50_ values in the range of 16.44 – 38.18 μM as compared to standard drug doxorubicin, IC_50_ = 0.34 ± 0.01, 1.6 ± 0.06, 0.92 ± 0.01, and 0.5 ± 0.07 μM against AU565, BT-474, MCF-7, and MDA-MB-231 breast cancer cell lines, respectively.

**Table 2:**
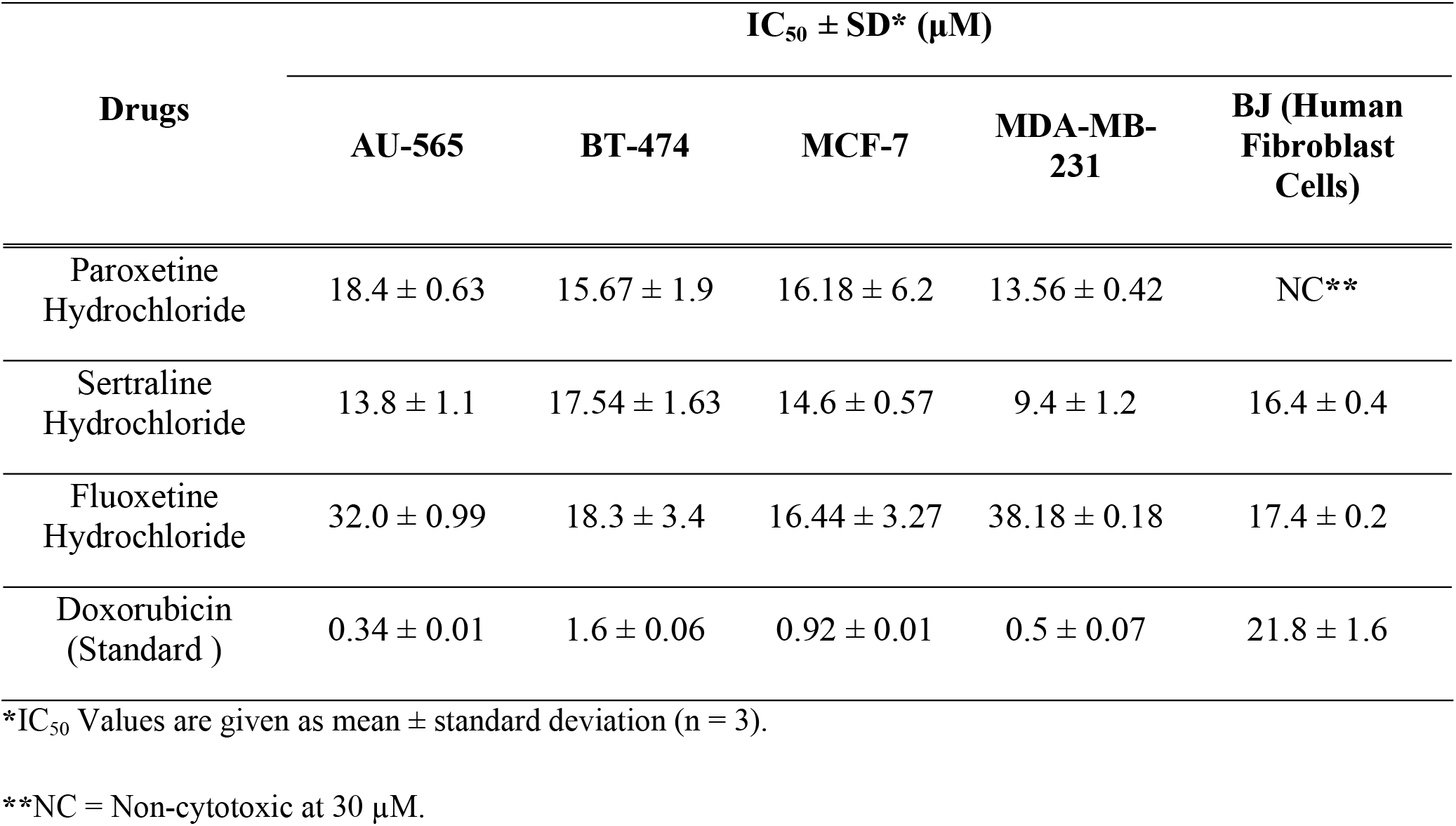
IC_50_ Values of anti-depressant drugs against breast cancer, and fibroblast cell lines.

| Drugs | $IC_{50} \pm SD^* (\mu M)$ | | | | |
| --- | --- | --- | --- | --- | --- |
|  | AU-565 | BT-474 | MCF-7 | MDA-MB-231 | BJ (Human Fibroblast Cells) |
| Paroxetine Hydrochloride | $18.4 \pm 0.63$ | $15.67 \pm 1.9$ | $16.18 \pm 6.2$ | $13.56 \pm 0.42$ | NC** |
| Sertraline Hydrochloride | $13.8 \pm 1.1$ | $17.54 \pm 1.63$ | $14.6 \pm 0.57$ | $9.4 \pm 1.2$ | $16.4 \pm 0.4$ |
| Fluoxetine Hydrochloride | $32.0 \pm 0.99$ | $18.3 \pm 3.4$ | $16.44 \pm 3.27$ | $38.18 \pm 0.18$ | $17.4 \pm 0.2$ |
| Doxorubicin (Standard ) | $0.34 \pm 0.01$ | $1.6 \pm 0.06$ | $0.92 \pm 0.01$ | $0.5 \pm 0.07$ | $21.8 \pm 1.6$ |

\* $IC_{50}$ Values are given as mean ± standard deviation (n = 3).
\*\*NC = Non-cytotoxic at $30 \mu M$ .

Cytotoxic potential of these drugs was also evaluated on BJ human fibroblast cell line. Paroxetine was found non-cytotoxic against human fibroblast cells. Whereas, fluoxetine hydrochloride (IC_50_ = 17.4 ± 0.2 μM), and sertraline hydrochloride (IC_50_ = 16.4 ± 0.4 μM) showed a comparable cytotoxicity to standard drug, doxorubicin (IC_50_ = 21.8 ± 1.6 μM).

As sertraline hydrochloride showed the most significant inhibition of all the breast cancer cell lines, we choose this drug for detailed study. Anticancer activity of sertraline hydrochloride has already been reported on MCF-7, and MDA-MB-231 breast cancer cell line [24, 25]. Therefore, consecutive studies were performed on HER2+ AU565 breast cancer cell line.

### 3.2 Sertraline Hydrochloride Induces Apoptosis in AU565 Breast Cancer Cells

Figs 2A, and B show that the percentage of live cells decreased significantly (p< 0.001) following the treatment with sertraline hydrochloride at all concentrations (7, 14, and 28 μM). Annexin/PI staining showed an increase in the percentage of early, and late apoptotic cells after treatment with 7, and 14 μM of sertraline hydrochloride. Maximum number of cells (34.34%) undergoing apoptosis were observed at 14 μM, while a decrease was observed at 28 μM. Necrotic cell death *i.e*. cells stained with PI only, was observed at higher concentration (28 μM) of sertraline hydrochloride.

**Fig. 2.**
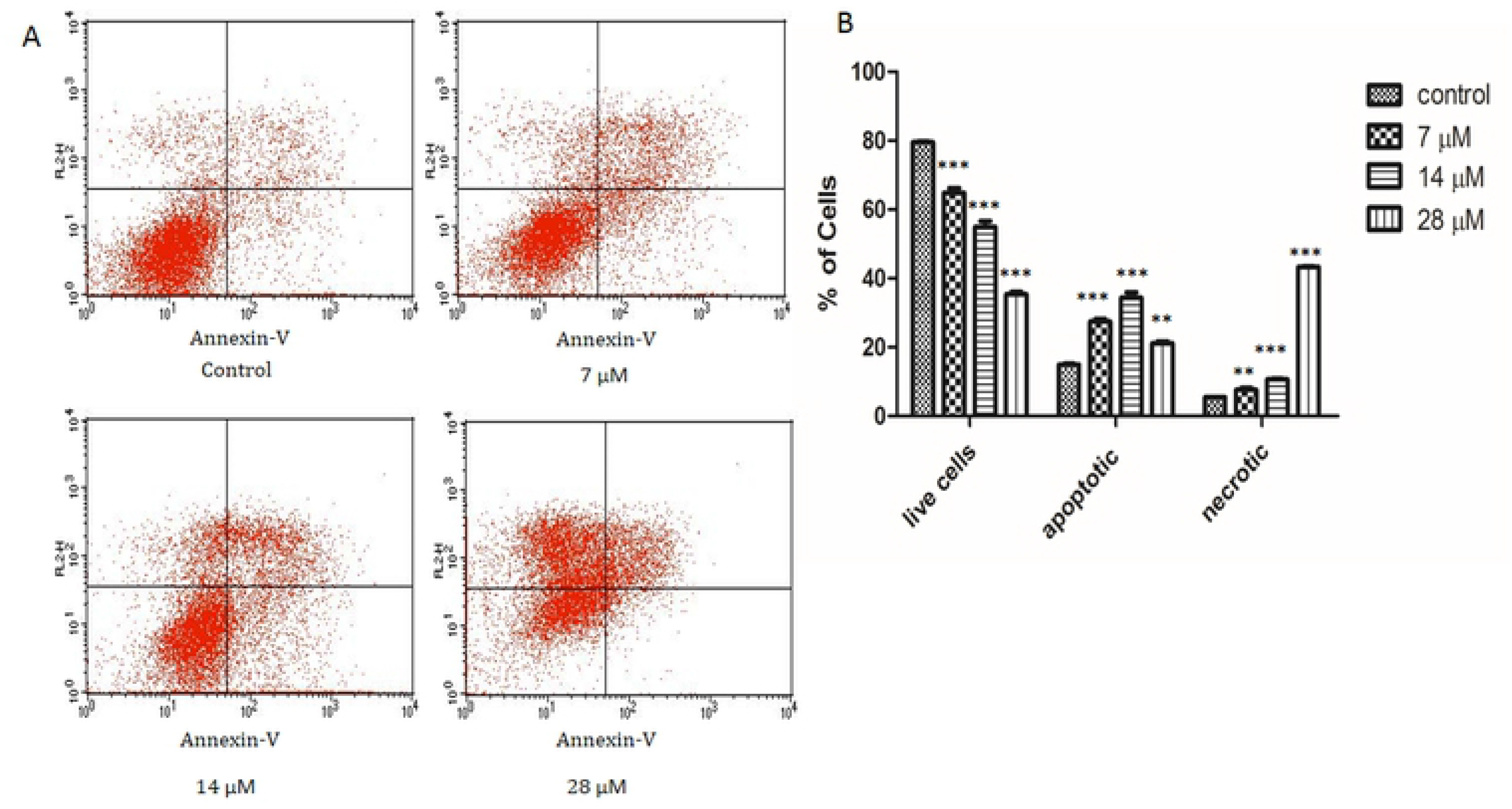
Sertraline hydrochloride induced apoptosis, in AU565 cells. (A) Annexin-V/PI dual staining assay using flow cytometry. Cells stained with annexin and/or PI are shown in different quadrants. Lower left (live cells); Lower right (early apoptotic cells); Upper right (late apoptotic) cells; Upper left (necrotic cells). (B) Quantification of cells sorted by dual staining of annexin V-FITC/PI. Results are expressed as mean ± SD (n=3). Significant differences are represented by **p < 0.01, ***p < 0.001

DNA fragmentation assay (DAPI) also revealed nuclear shrinkage, and DNA fragmentation followed by the treatment with different concentrations (7-28 μM) of sertraline hydrochloride, as shown in Fig 3. In accord with our results of flow cytometry, decrease in live cells a view of debris was observed after treatment with 28 μM of sertraline hydrochloride, which further suggests necrotic cell death at higher concentration.

**Fig 3.**
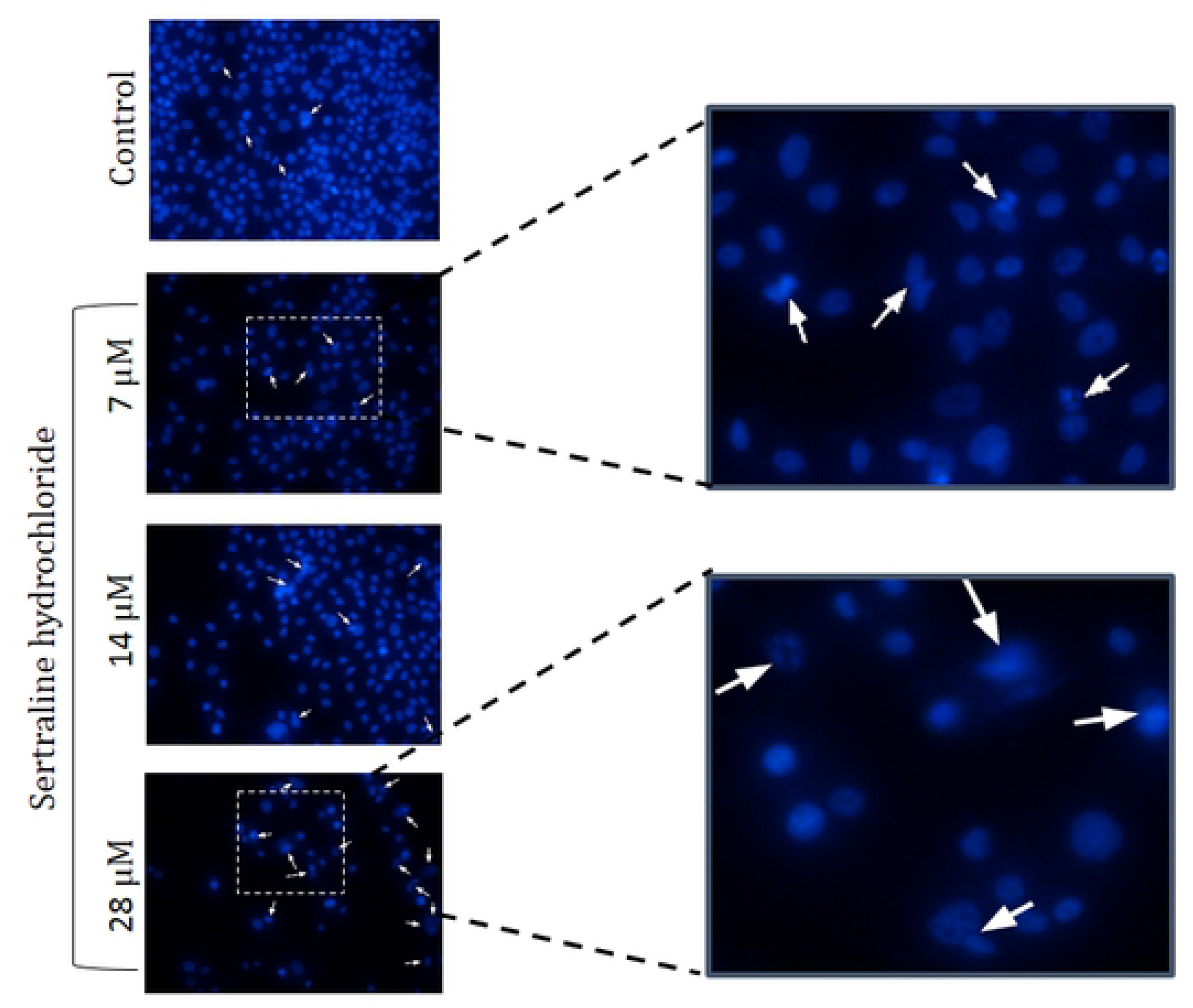
Sertraline hydrochloride caused nuclear fragmentation. Visualization of DNA damage in response to sertraline hydrochloride by DAPI staining, images were captured at 20x, 25 μm.

### 3.3 Sertraline Hydrochloride Activates Executioner Caspase-3/7

Since nuclear condensation, DNA fragmentation, and increase in Annexin (+) cells, are characteristic indicative of apoptotic cell death. It was important to ascertain the involvement of caspase, which is a hallmark of apoptosis [26]. Caspase-3/7 activity was monitored at different time interval. After 6 h of treatment caspase activity was comparable to untreated cells (0.97, and 1.5 fold at 7, and 14 μM, respectively). Caspase activity peaked between 12–24 h, and a significant increase (p <0.01, 0.001) was observed, as compared to untreated cells. Maximum caspase-3/7 activity was observed after 24 h of treatment (4.9 fold at 14 μM) (Fig. 4). Decrease in caspase activity was observed after 48 h at both the concentrations used (7, and 14 μM).

**Figure 4.**
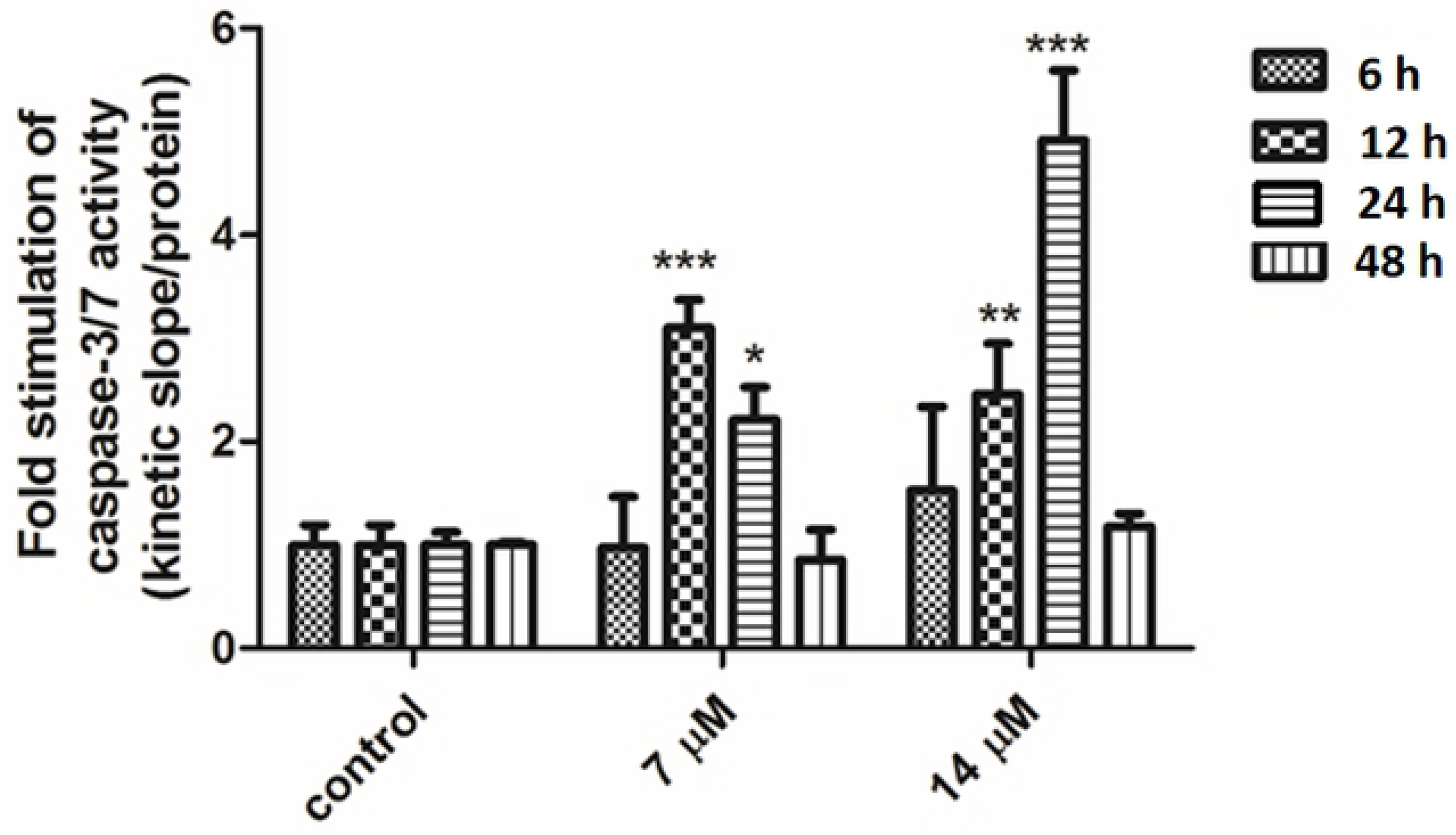
Caspase-3/7 activity increases after treatment with sertraline hydrochloride. Relative caspase-3/7 activity at 6, 12, 24, and 48 h of treatment. Results are expressed as mean ± SD (n=3). Significant differences are represented by *p < 0.05, **p < 0.01, ***p < 0.001.

### 3.5 Sertraline Hydrochloride arrest Cells at G2/M Phase of Cell Cycle

Cell cycle analysis by flow cytometry (Figs 5A, and B) revealed that sertraline hydrochloride mediates its cytotoxic effect through perturbation of cell cycle. Sertraline hydrochloride causes a significant decrease (p< 0.05, <0.01, and <0.001) in G0/G1 (growth) phase of AU565 cells as compared to untreated cells (28.1–44.7 *vs* 58.6% in control). A significant (p < 0.05, and p < 0.01), and dose-dependent decrease in DNA synthesis (S) phase following treatment with sertraline hydrochloride (21.6, 20.8, and 18.5% cells at 7, 14, and 28 μM, respectively, *vs* 41.7% cells in control), accompanied by accumulation of cells in the mitotic phase (G2/M) phase (36.0 – 49.7%) was also observed. Maximum G2/M phase arrest was observed at 7 μM (49.7 *vs* 3.657%; p < 0.001), as compared to untreated cells. These results indicated that sertraline drives mitochondrial cell death in AU565 breast cancer cells.

**Fig 5.**
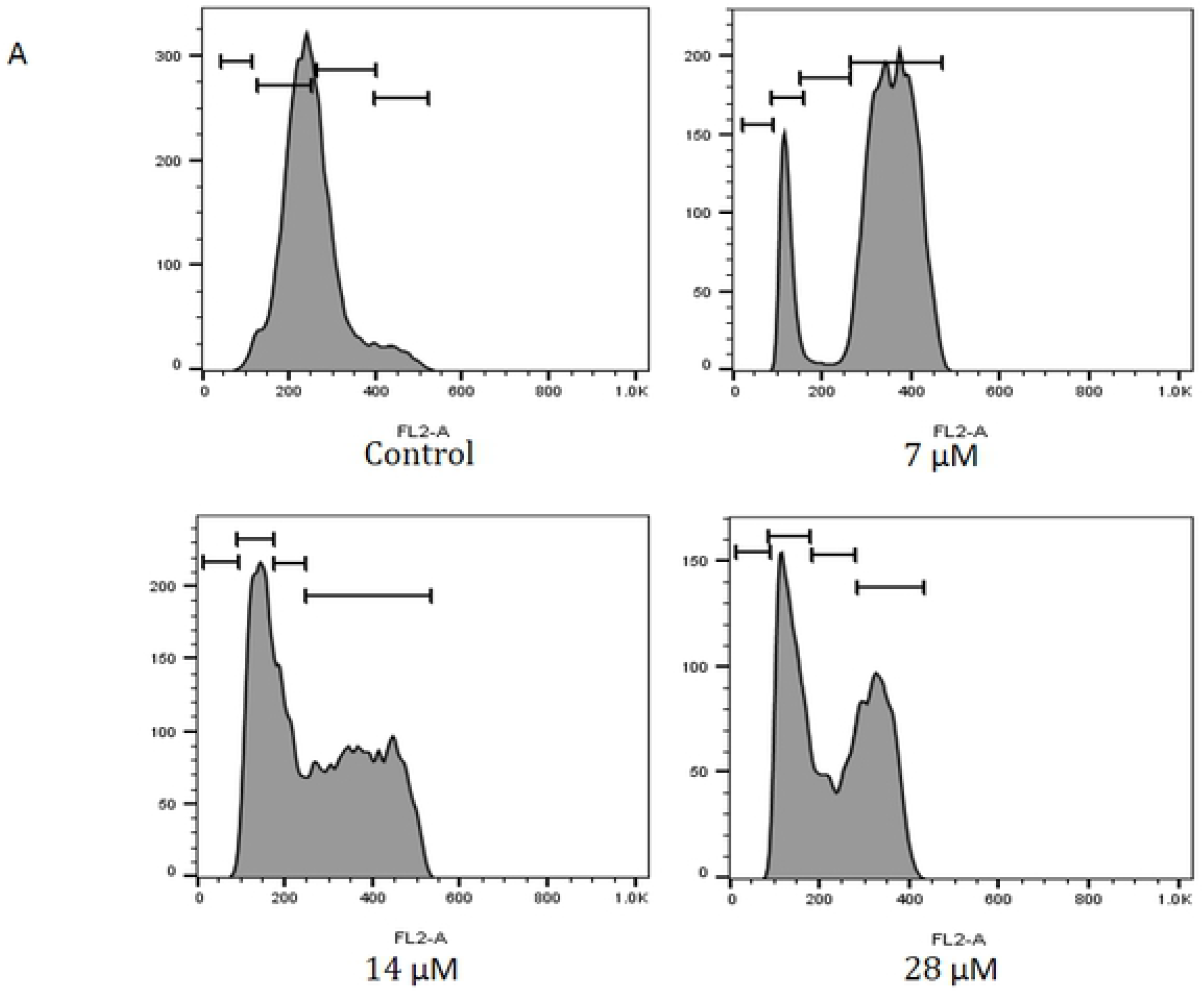
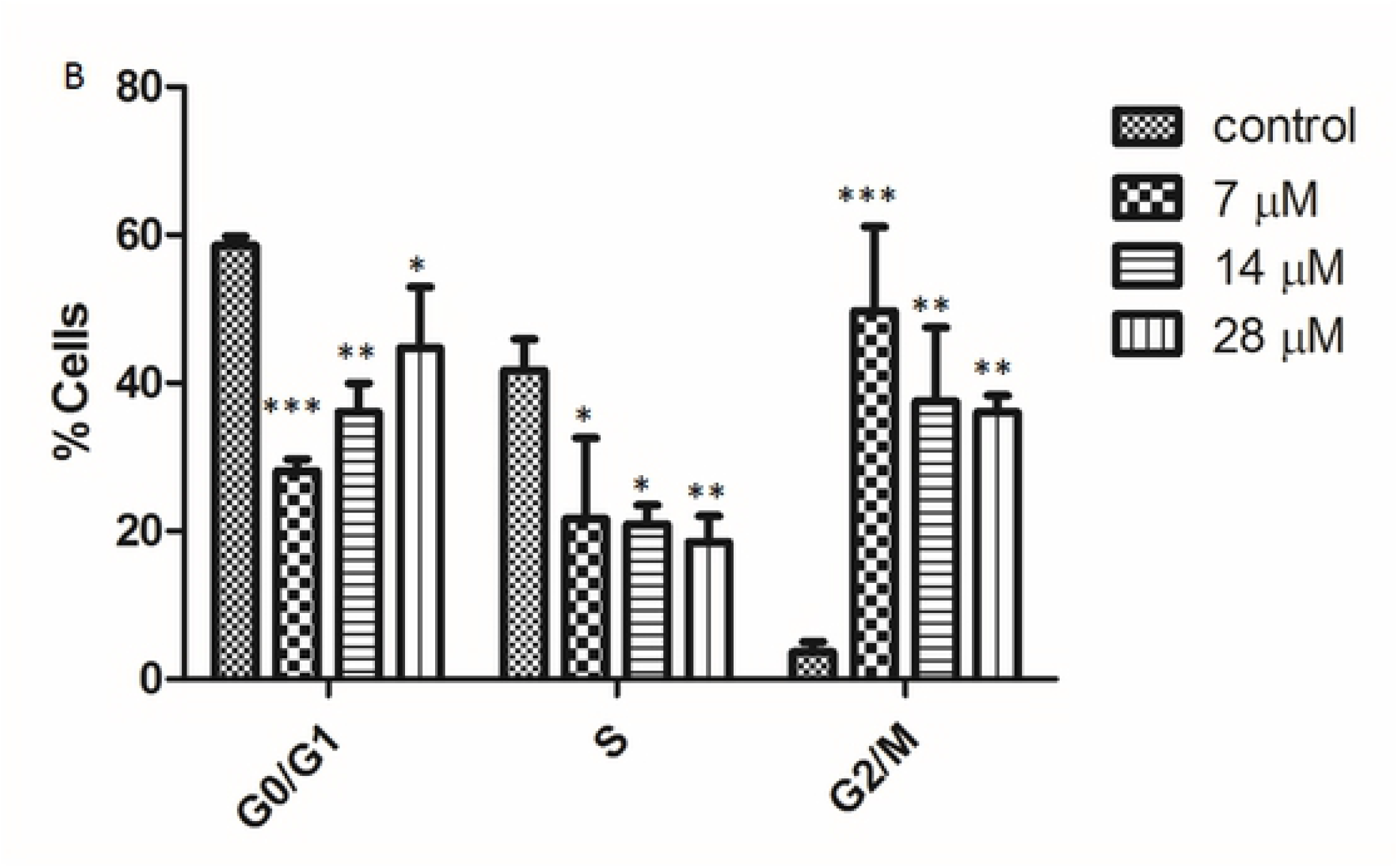
Sertraline hydrochloride caused a G2/M cell cycle arrest. (A) Flow cytometric analysis of cells treated with different concentrations of sertraline hydrochloride for 48 h. (B) Quantification of cells distributed in different phases of cell cycle progression, relative cell population ± SD (n=3). Significant differences are represented by *p < 0.05, **p < 0.01, ***p < 0.001.

### 3.6 Expression of Pro/Anti-apoptotic Genes in Sertraline-Treated AU565 Cells

Apoptosis is governed by the expression of certain pro-, and anti-apoptotic genes. An insignificant increase (1.58 fold) in the expression of pro-apoptotic Bax gene was observed, whereas BCl2 gene (anti-apoptotic) showed a comparable expression (0.97 fold) at 7 μM to untreated cells. While, c-myc oncogene showed comparable expression (0.8 fold) after treatment of AU-565 cells with 14 μM of sertraline hydrochloride (Fig 6). Hence it could be proposed that sertraline hydrochloride mediates BCl2-independent cell death in AU565 breast cancer cells.

**Fig 6.**
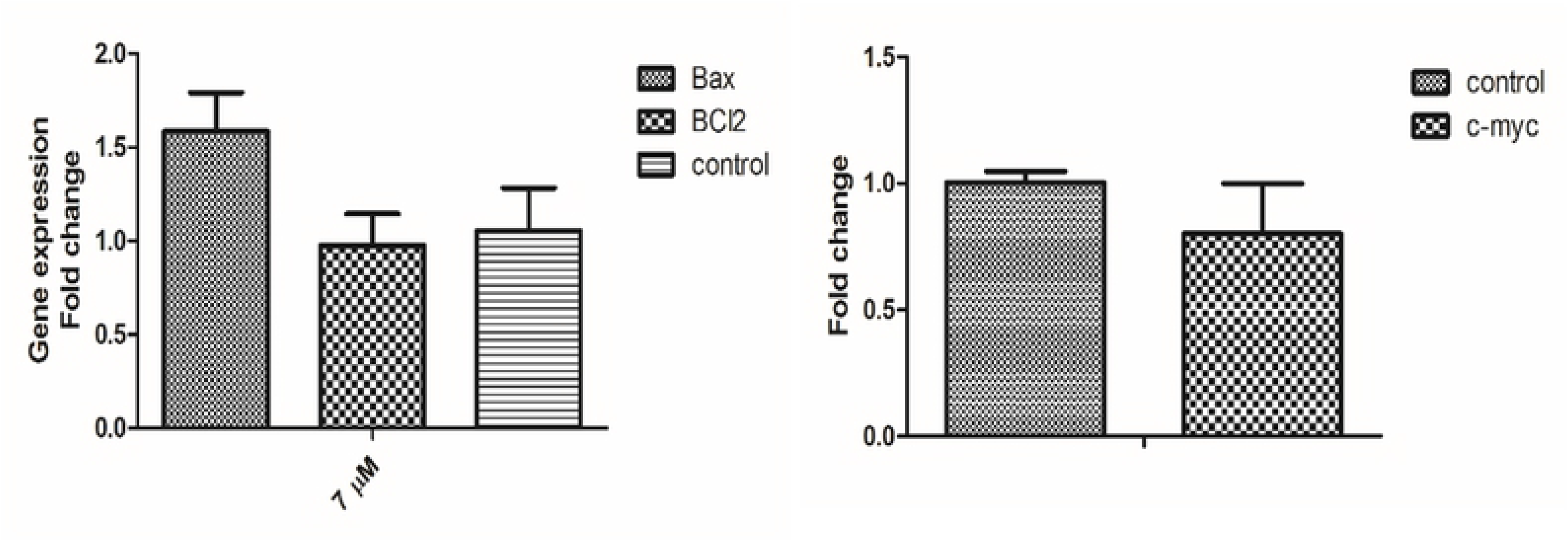
Expression of pro-, and anti-apoptotic genes in the presence of sertraline hydrochloride. Expression of Bax, and BCl2 (left, 7 μM), c-myc oncogenes (right, 14 μM). Results are expressed as mean ± SD (n=3).

## 4. Discussion

The concept of drug repositioning has recently drawn attention for drug discovery and development. This approach is widely used, and has been successful in several cases e.g., aspirin as an anti-platelet medication, sildenafil for erectile dysfunction [27, 28]. To complement the ongoing efforts of developing better chemo-therapeutic agents, the effect of different anti-depressants (Table 2) on viability of MCF-7, MDA-MB-231, BT-474, and AU565 breast cancer cell lines was assessed during the present study.

Sertraline hydrochloride is one of the most commonly used serotonin inhibitor. It is well established that depression could assist the on-set breast cancer, and certain anti-depressants, particularly SSRIs has been reported to possess anti-cancer effect. But the effect of these drugs on HER2+ cancers remain scarce. Results of our study provides convincing evidences for the anti-cancer effect of sertraline hydrochloride against HER2+ AU565 cell line. Nuclear shrinkage, fragmentation, and DNA damage are important indications of mitochondrial cell death. Sertraline-induced apoptosis was inferred from flow cytometry and DAPI staining (Fig. 3). Apoptotic effect of sertraline has been reported earlier in hepatic cells [29]. Hodge *et al.* also indicated that apoptosis is responsible for the decreased cell viability in sertraline- and paroxetine-treated human osteoclasts, and osteoblast cells [30]. Exposure of cells to different concentrations of sertraline hydrochloride caused nuclear shrinkage, and extensive DNA fragmentation, as compared to untreated cells. Apoptotic bodies were observed after treatment with 28 μM of sertraline hydrochloride. Moreover, only few cells were found alive, and a view cell debris was observed, which suggested the combination of necrotic, and apoptotic cell death. Our results are in agreement with the previous studies which showed that SSRIs can increase the DNA damage [31, 32]. It is reported that SSRIs result in misreading of the DNA code by inhibiting the DNA binding to apetala 2 (AP-2), and thus cause DNA damage [33]. DNA damage was further confirmed through arrest of cell cycle. Previous studies indicated that SSRIs induced apoptosis is accompanied by population arrest in different phases of cell cycle. Gil-Ad and co-workers also showed that sertraline induced a significant decrease in synthetic phase (S-phase) of the cell cycle, and arrest colon cancer cells at G0/G1 phase [34]. In order to study the effect of sertraline hydrochloride on cell cycle progression of AU565 breast cancer cells, distribution of cells in different cell cycle phases was studied using flow cytometry. G2/M phase arrest was observed at lower concentration (7 μM), while higher concentrations of sertraline hydrochloride (14, and 28 μM) caused cells to exit from G2/M phase, indicating a possible induction of apoptosis, leading to cell death. Sertraline-induced G2/M phase arrest is also evident in acute myeloid leukemia cells [35]. It is also reported that fluoxetine hydrochloride exerted its anti-proliferative action by suppressing cell cycle progression in breast cancer, neuroblastoma, and colon cancer cell lines [36]. Time dependent increase of executioner caspase-3/7 was also observed up to 24 h incubation of AU-565 breast cancer cells with sertraline hydrochloride. These results suggest that sertraline hydrochloride activates the caspases to induce DNA fragmentation, and apoptosis. Decrease in caspase-3/7 activity after 48-h could be attributed to further cleavage of caspase-3, and cell death. Consistent results of sertraline-induced mitochondrial dysfunction, and cell death was observed in astrocytes, and sertraline-associated hepatotoxicity [37, 38]. Thus activation of caspase-3/7 after treatment with sertraline hydrochloride further corroborated our results and demonstrate that sertraline hydrochloride mediated cytotoxicity AU565 cells is a result of apoptotic cell death. Since caspase-3 is an executioner which is activated in both intrinsic and extrinsic cell death, further studies are warranted to reveal the molecular mechanisms involved in anti-cancer effects of sertraline hydrochloride.

Results of gene expression through PCR showed only marginal insignificant increase in apoptotic Bax, while the expression of BCl2, and oncogene c-myc remains unchanged. This is an unusual finding compared with the observations made with other apoptotic stimuli. Although it has been shown that, anti-tumor drugs, such as flavopiridol can induce apoptosis through different pathways of caspase activation which is largely independent of Bcl-2 [39]. Based on these findings we can suggest that activation of caspases are instrumental in sertraline-induced apoptosis. Overall our study indicate that cell death elicited by sertraline hydrochloride in AU565 breast cancer cells involves apoptotic pathways as evidenced by nuclear shrinkage, DNA fragmentation, and Annexin (+) cells. Necrotic cell death was observed at higher concentrations, which certainly merits further investigations.

## 5. Conclusion

To the present, this is the first study to illustrate the possible mechanism of cytotoxicity of sertraline hydrochloride on AU565 breast cancer cells. Results of the present study has shown that anti-depressant sertraline hydrochloride causes the arrest of cell cycle, nuclear fragmentation, and caspase-3 activation in AU565 cells. These results suggest that sertraline hydrochloride induced cell death in AU565 breast cancer cells possibly through caspase dependent apoptotic pathway. More studies are warranted to investigate the underlying mechanisms, which may provide a key to elucidate sertraline’s unusual mechanism of action.

## Acknowledgement

ATW and MIC acknowledge the enabling role of the Higher Education Commission, Pakistan, through a financial support under National Research Programme for Universities (NRPU) # 20-3790 entitled, “Studies on the Chemoprevention of Mammary Carcinogenesis by Dietary Agents”.

## Author Contributions

Conceptualization: M. Iqbal Choudhary, and Atta-ur-Rahman

Formal analysis: Atia tul-Wahab, Rafat A. Siddiqui

Methodology: Sharmeen Fayyaz, Rimsha Irshad

Project administration: Atia-tul-Wahab, M. Iqbal Choudhary, Atta-ur-Rahman

Supervision: Atia tul-Wahab, M. Iqbal Choudhary

Writing – original draft: Sharmeen Fayyaz

Writing – review & editing: Atia tul-Wahab, Rafat A. Siddiqui, M. Iqbal Choudhary.

